# Auditory steady-state responses: multiplexed amplitude modulation frequencies to reduce recording time

**DOI:** 10.1101/2023.06.01.543200

**Authors:** Rien Sonck, Jonas Vanthornhout, Estelle Bonin, Tom Francart

## Abstract

**Objectives:** This study investigated the efficiency of a multiplexed amplitude-modulated (AM) stimulus in eliciting auditory steady-state responses (ASSR). The multiplexed AM stimulus was created by simultaneously modulating speech-shaped noise with three frequencies chosen to elicit different neural generators: 3.1, 40.1, and 102.1 Hz. For comparison, a single AM stimulus was created for each of these frequencies, resulting in three single AM conditions and one multiplex AM condition.

**Design:** Twenty-two bilaterally normal-hearing participants (18 female) listened for 8 minutes to each type of stimuli. The analysis compared the signal-to-noise ratios (SNRs) and amplitudes of the evoked responses to the single and multiplexed conditions.

**Results:** The results revealed that the SNRs elicited by single AM conditions were, on average, 1.61 dB higher than those evoked by the multiplexed AM condition (p < 0.05). The single conditions consistently produced a significantly higher SNR when examining various stimulus durations ranging from 1 to 8 minutes. Despite these SNR differences, the frequency spectrum was very similar across and within subjects. Additionally, the sensor space patterns across the scalp demonstrated similar trends between the single and multiplexed stimuli for both SNR and amplitudes. Both the single and multiplexed conditions evoked significant ASSRs within subjects. On average, the multiplexed AM stimulus took 31 minutes for the lower bound of the 95% prediction interval to cross the significance threshold across all three frequencies. In contrast, the single AM stimuli took 45 minutes and 42 seconds.

**Conclusion:** These findings show that the multiplexed AM stimulus is a promising method to reduce the recording time when simultaneously obtaining information from various neural generators.

## Introduction

The auditory steady-state response (ASSR) is a commonly used objective measure of the auditory system (Picton et al., 2003; Picton & Burkard, 2012). During the test, electrodes are placed on the scalp to record the electrical signal generated by the brain in response to an auditory stimulus, requiring minimal subject participation. The ASSR uses an amplitude-modulated (AM) carrier wave. Listening to this stimulus evokes sustained neural activity that is phase-locked to the stimulus’s modulation frequency. The response can be elegantly and fully automatically detected by signal processing and statistical methods determining when the modulation frequency is significantly present in the electroencephalogram (EEG) (John & Picton, 2000; Van Dun et al., 2008).

The stimulus used to evoke ASSRs is characterized by two important frequencies: the carrier frequency (or frequency range), which determines which part of the cochlea is activated, and the modulation frequency, which relates to the main neural generators of the recorded responses (Herdman et al., 2002). Multiple carrier frequencies, combined with multiple modulation frequencies that are very close together, have been used to simultaneously and (more or less) independently assess multiple areas of the cochlea (i.e., multiple ASSR; John et al., 1998; Lins et al., 1994, 1996). The carrier and modulation frequencies can be independently controlled, and stimuli containing multiple simultaneous frequencies have been deployed to decrease overall recording time (Luts & Wouters, 2004).

In multiple ASSRs, the modulation frequencies are usually selected from a small range primarily targeting one neural generator, such as 30 to 50 Hz (e.g., Gransier et al., 2017; John et al., 1998) or 70 to 110 Hz (e.g., Dimitrijevic et al., 2002; Hatton & Stapells, 2010; Holmes et al., 2018; Lins et al., 1994, 1996; Luts et al., 2006). In contrast, we will focus on multiple simultaneous modulation frequencies *that are far apart*, targeting multiple neural generators. The cortical areas primarily respond to low AM frequencies (< 20 Hz; Alaerts et al., 2009; Farahani et al., 2020; Herdman et al., 2002; Y. Wang et al., 2012), while subcortical areas such as the brainstem primarily respond to higher AM frequencies (> 80-100 Hz; Bidelman, 2015; Farahani et al., 2020; Herdman et al., 2002; Luke et al., 2017).

Other stimuli, like the multi-tone frequency (Dolphin, 1996, 1997), have explored a wider frequency range, but these did not include frequencies below 30 Hz and were exclusively tested in dolphins. Other studies have explored the use of complex tones emphasizing fundamental frequencies, speech syllable envelopes, and prosodic fluctuations. However, these studies also predominantly target specific neural generators, either the brainstem or cortical areas, without encompassing the entire auditory pathway (Henry et al., 2014; Varghese et al., 2015; L. Wang et al., 2019).

In contrast, our objective is to develop a novel type of ASSR, termed the “multiplexed AM stimulus,” capable of simultaneously and efficiently gathering information from various neural generators within the auditory system. The proposed multiplexed stimulus involves simultaneously amplitude-modulating the carrier wave with multiple frequencies selected from a broad frequency range (< 10 Hz to > 100 Hz). This broad range allows the stimulus to simultaneously target multiple neural generators in the auditory pathway, such as the brainstem, midbrain, and auditory cortex. Consequently, the multiplexed stimulus has the potential to efficiently extract information from various neural generators, reducing recording time and reducing the impact of fatigue and loss of attention.

## Materials and Methods

### Participants

We recruited 22 bilaterally normal-hearing subjects (4 male and 18 female) with a mean age of 22.7 ± 2.6 years. We conducted pure-tone audiometry on 15 participants to assess their hearing thresholds across the frequency range of 125 Hz to 8 kHz, with the threshold for normal hearing set at <20 dB HL. The remaining participants reported no issues with their hearing. All subjects participated voluntarily and signed an informed consent. The Ethics Committee Research UZ / KU Leuven approved the experiment (S57102).

### Experimental procedure

At the start of the experiment, subjects were placed in a soundproof Faraday shielded booth, and ASSR stimuli were binaurally presented via ER-3A insert earphones (Etymotic Research Inc, IL, USA) to each subject. The earphones were encased in an electrically grounded box. Subjects partook in three single amplitude-modulated conditions and one multiplexed AM condition, lasting around 32 minutes. During each condition, subjects listened for eight minutes to an amplitude-modulated carrier presented using the APEX software (Francart et al., 2008). The order of these conditions was randomized for each subject. The stimulus was presented using an RME Fireface UC soundcard (Haimhausen, Bayern, Germany) at 80 dBA.

### Stimulus construction

We employed speech-shaped noise to generate the stimulus carrier, see Figure 1A, crafted by extracting the spectrum from a self-recorded audiobook with a male speaker that is not publicly accessible online, as it was exclusively recorded within our laboratory. The choice of the audiobook stems from its use in a broader research project. This noise carrier was 100% modulated in the single AM conditions by either a 3.1 Hz, a 40.1 Hz, or a 102.1 Hz sinusoidal modulator. The modulator varies between -1 and 1 in this technique, as depicted in Figure 1B. Due to this, the modulation frequency is artificially double as such, when targeting a specific frequency, *f*, in the modulation process, we divide by two when creating the sinusoidal modulator *m*_*f*_, as illustrated in Equation 1, where *t* stands for time. This modulator will ultimately stimulate frequency *f* in the modulation spectrum, see Figures B and C. The stimulus signal is created by multiplying the carrier with the modulator.

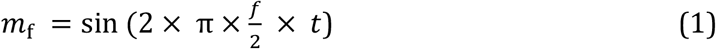

We chose these frequencies based on the neural generators they target. Specifically, the 3.1 Hz AM frequency targets the cortical areas, the 102.1 Hz AM frequency targets the brainstem, and the 40.1 Hz AM frequency targets the cortical areas, the brainstem, and the thalamus. For the multiplexed AM condition, the carrier was simultaneously 100% modulated by all AM frequencies, i.e., a multiplexed modulator; see Equation 2, where *m*, the multiplexed modulator, is the element-wise multiplication of the 3.1 Hz sinusoidal modulator (*m*_3_), the 40.1 Hz sinusoidal modulator (*m*_40_) and the 102.1 Hz sinusoidal modulator (*m*_102_). From here on, we will refer to the stimuli and evoked potentials as 3 Hz, 40 Hz, and 102 Hz for convenience.

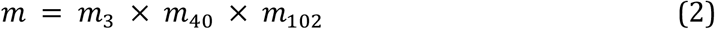

Figure 1B-C shows the multiplexed modulator and the resulting multiplexed stimulus. An amplitude reduction is introduced in the resulting signal by multiplying three single sinusoidal modulators to create a multiplexed modulator.

**Figure 1.**
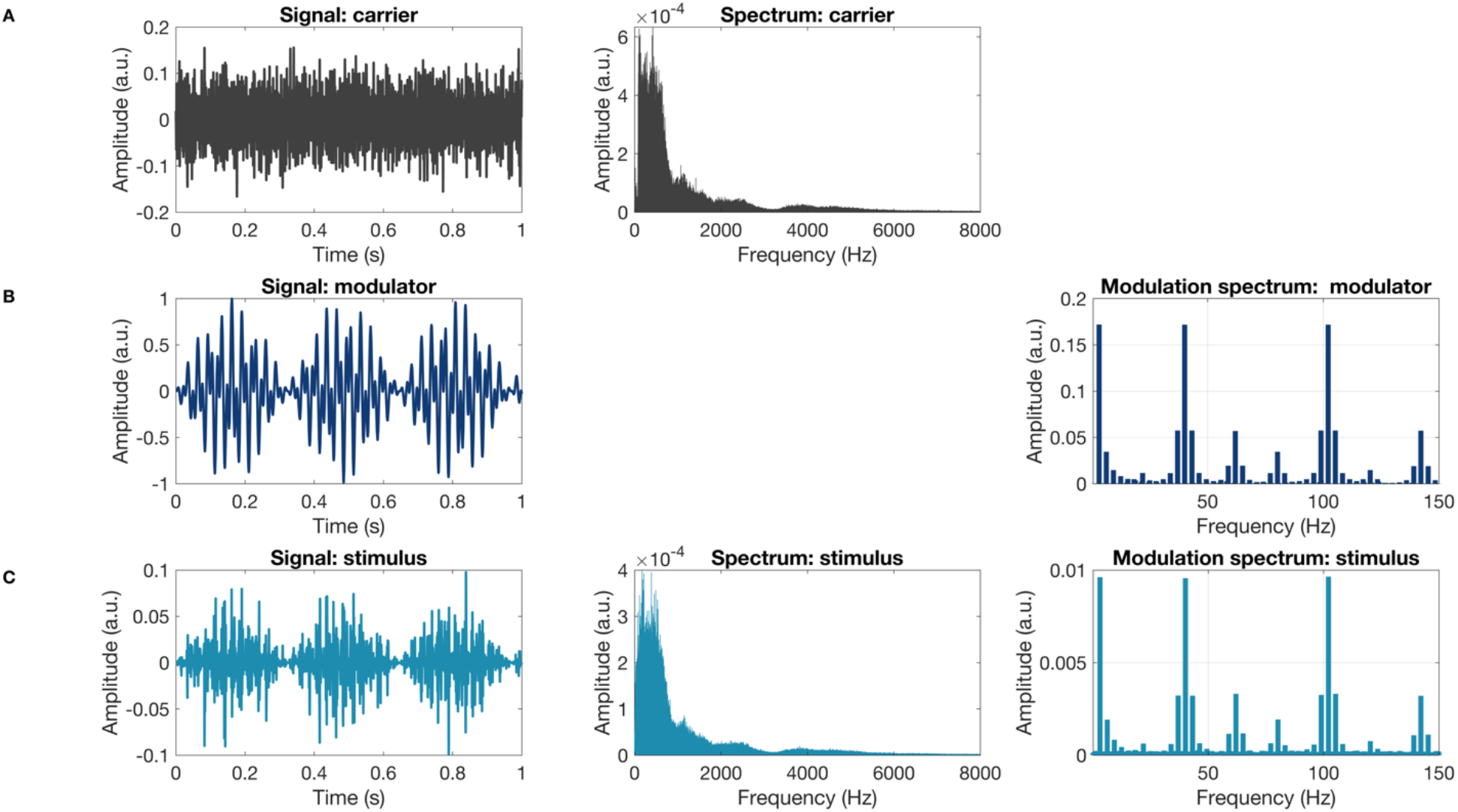
(A) Depicts the speech-shaped noise carrier and its corresponding spectrum. (B) Illustrates the modulator and its modulation spectrum, while (C) showcases the resulting multiplexed signal (on the left), including its spectrum (in the middle) and modulation spectrum (on the right). The modulation spectrum shows the stimulus spectrum after passing through the hair-cell non-linearity by using full-wave rectification.

We assessed both the time and frequency domain to quantify this attenuation for each AM frequency. In the time domain, we calculated the Amplitude Attenuation Index (AAI); see Equation 3.

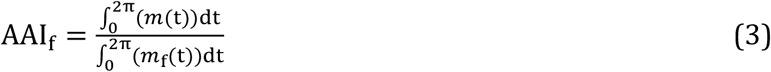

In the equation, the multiplexed modulator is denoted as *m*(*t*), where *t* stands for time, and *m*_*f*_(*t*) represents one of the single sinusoidal modulators used to create the multiplexed modulator. The *AAI*_*f*_ value indicates the amplitude attenuation experienced by a specific frequency component *f* within the multiplexed modulator. An *AAI*_*f*_ of 1 signifies that the frequency component in the multiplexed modulator did not undergo any amplitude attenuation. Conversely, an *AAI*_*f*_ of 0 suggests a complete amplitude attenuation of the frequency component within the multiplexed modulator. The *AAI* for each frequency component in the multiplexed modulator is shown in Figure 2. Since the frequencies 3 Hz, 40 Hz, and 102 Hz have no common denominator, the *AAI* varies across different periods of the multiplexed modulator. However, on average, the *AAI* is 0.4 for each frequency component, which indicates that the amplitude of each frequency component is only 0.4 of its original amplitude. Upon examining the frequency domain of both the multiplexed modulator and multiplexed stimulus, as illustrated in Figure 1B-C, both exhibit consistent amplitudes across the AM frequencies. Thus, the AM frequencies in the multiplexed signal experience amplitude attenuation compared to the single ASSR; however, the amplitudes between AM frequencies remain consistent.

**Figure 2.**
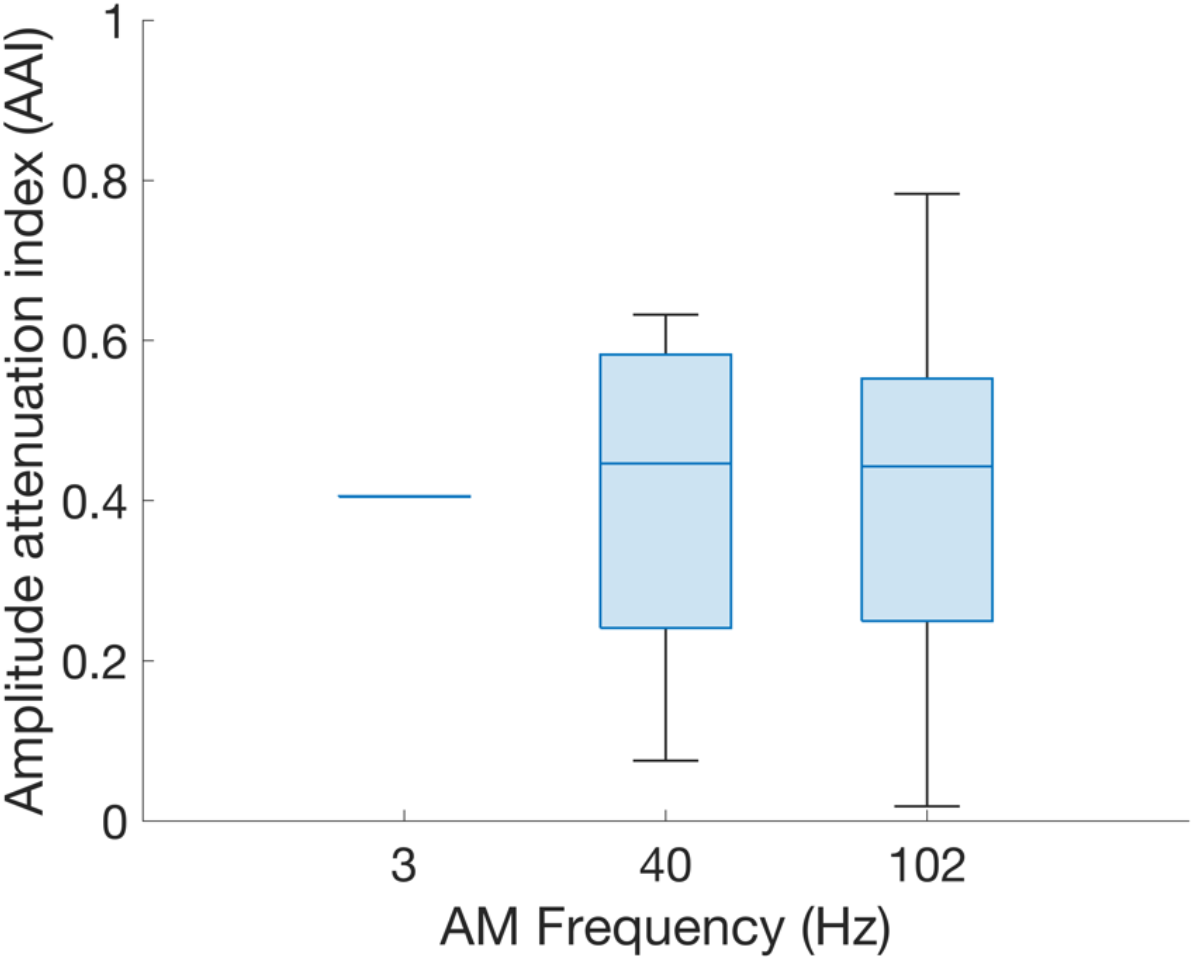
Shows the Amplitude Attenuation Index (AAI) corresponding to each amplitude-modulation frequency of the multiplexed modulator.

### EEG acquisition

Subjects’ brain activity was recorded using electroencephalography (EEG) with a 64-channel BioSemi ActiveTwo system (BioSemi, Amsterdam, Netherlands) with Ag/AgCl electrodes mounted on a cap according to the 10-20 system of electrode placement (Oostenveld & Praamstra, 2001). The EEG data were acquired using the ActiView software at an 8192 Hz sampling rate.

### Signal preprocessing

The electroencephalography (EEG) data preprocessing was performed offline in the MATLAB environment (“MATLAB” 2021). We extracted the amplitude of each frequency in the EEG signal by resampling the raw EEG to a sampling rate of 1024 Hz and applying a 20th-order infinite impulse response comb-notch filter to remove the 50 Hz powerline and its harmonics. Bad channels were identified using the root mean (RMS) method, which involves calculating the RMS value for each channel and comparing it to the threshold of 3 × the median RMS across channels. Channels with RMS values above this threshold are considered bad channels and were then interpolated using adjacent channels. The signal was then filtered using a 4th-order Butterworth high-pass zero-phase filter with a cutoff frequency of 1 Hz to remove DC and drift components. A multi-channel Wiener filter (Somers et al., 2018) was applied to remove eye-blinking artifacts. The signal was then re-referenced to the Cz electrode for all analyses, apart for the topographic maps, where average re-referencing was applied.

### Signal preprocessing for the SNR calculation

The same bad channels detected during the amplitude preprocessing were interpolated to extract the SNR of each frequency in the EEG signal. The signal was re-referenced to the Cz electrode and divided into epochs based on the stimulus frequency. Since the 3 Hz, 40 Hz, and 102 Hz do not have a common denominator to align their phases, we opted to make slight variations in epoch timing to ensure we have an integer number of cycles such that no phase drift could occur. For the 3.1 Hz stimulus, the signal was divided into 0.9677-second epochs, resulting in 7928 samples per epoch. For the 40.1 Hz and 102.1 Hz stimuli, the signal was divided into 0.9989-second epochs, resulting in 8184 samples per epoch. After dividing the signal into epochs, a quality control step was performed to identify and discard bad epochs. It was discarded if an epoch’s peak-to-peak value was higher than six times the median of the peak-to-peak values obtained across all the channels for each given epoch.

### Electrode selection

The SNR was first normalized within subjects and then averaged per modulation frequency using the min-max function, shown in Figure 3. Across modulation frequencies, the highest normalized SNR was observed for the P10 and P9 electrodes, located in the parietal-occipital region of the brain, see Figure 3B, and thus were chosen as our electrode selection. All results except for the topographic maps are averaged across P9 and P10.

**Figure 3.**
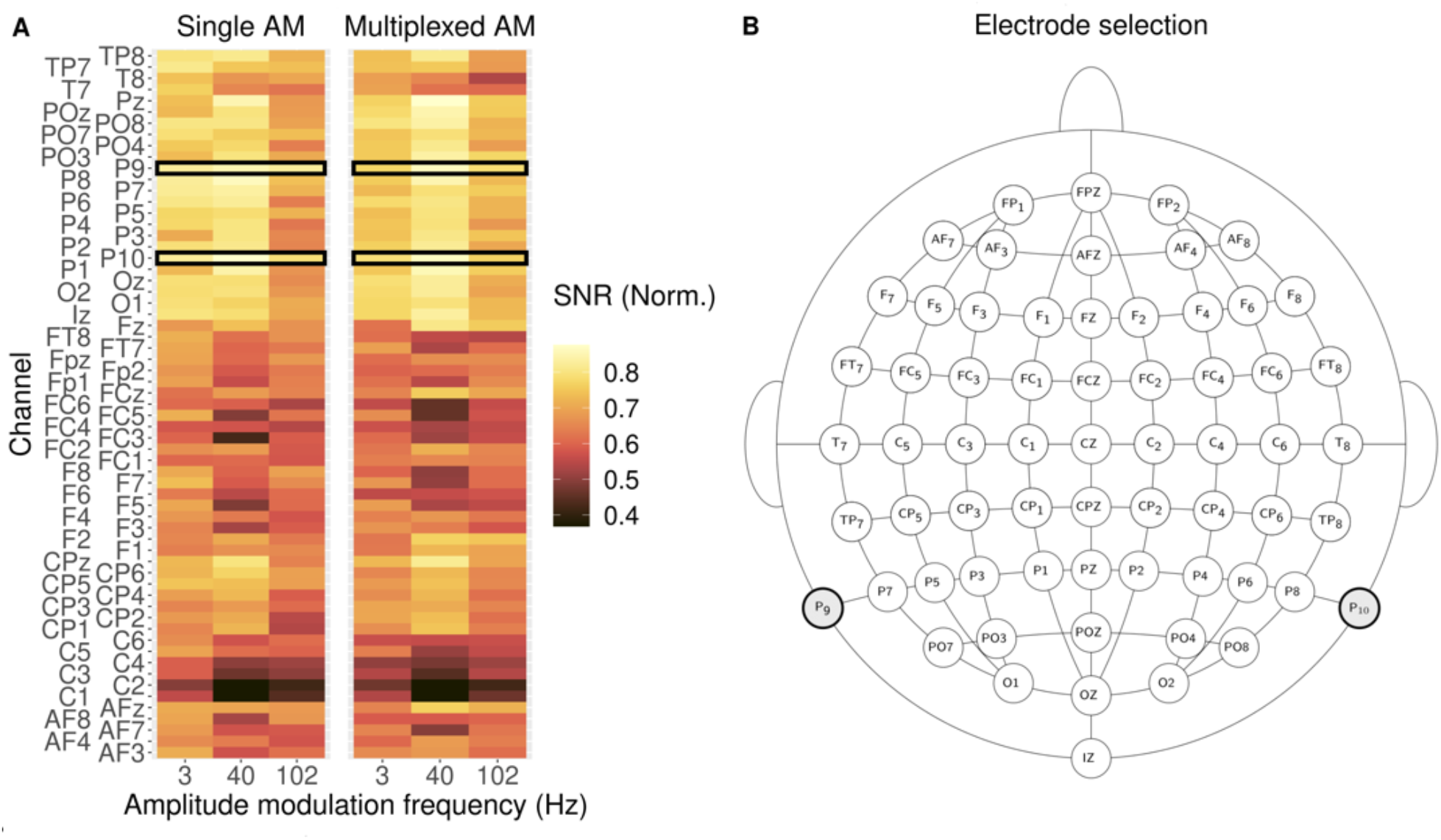
(A) The average signal-to-noise ratio (SNR) normalized per condition across participants. Electrode P10 and P9 show the highest SNR across modulation frequencies. (B) The electrode selection is shown on a 10-20 topographical electrode map.

### Outcome measures and statistics

#### Signal-to-noise ratio

Each frequency bin’s SNR was determined using Hotelling’s *T*^2^ statistic (Hotelling, 1992) a statistical approach that generalizes the student’s t-statistic to a multivariate dataset. In our study, we used Hotelling’s *T*^2^ statistic to compare the mean of the modulated frequency bin to the bin’s variability across epochs, allowing for the detection of synchronized activity in the EEG signal. We applied the Hotelling *T*^2^ statistic to each EEG channel to determine if the synchronized activity evoked by the ASSR stimuli was significantly different from the non-synchronized neural background activity.

#### Linear mixed effect model: single vs. multiplexed

We constructed a linear mixed-effect (LME) model using R (R Core Team 2020) and the lme4 package (Bates et al., 2015) to investigate the relationship between listening to a multiplexed AM stimulus, listening to a single AM stimulus, and the various AM frequencies on the SNR of the evoked response. Initially, we fit a linear mixed-effects (LME) model to our data, including all predictor variables and their interactions. However, the interaction effects between the AM frequencies and the ASSR condition (single vs. multiplexed) were found to be non-significant and were subsequently removed from the model, see Equation 4.

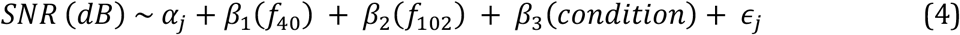

The model includes three categorical variables: *f*40 (representing a 40 Hz AM frequency), *f*102 (representing a 102 Hz AM frequency), and *condition* (representing the ASSR condition, with 0 indicating single and 1 indicating multiplexed). These variables are considered fixed effects in the model. The 3 Hz AM frequency is also included in the model’s intercept. Interaction terms are added between the AM frequencies and the conditions. As random effects, the intercepts for the subjects *α*_*j*_ are included to account for differences between subjects, and the residual error term *ϵ*_*j*_ represents the variability that the model does not explain. Although the residuals of our linear-mixed effect model do not follow a normal distribution, resulting in less model power, we still proceeded with the planned analysis. To interpret the LME model, we conducted post-hoc tests using the emmeans package in R (Lenth et al., 2023) with Tukey’s multiple comparison test (Keselman & Rogan, 1977).

#### Linear mixed effect model: SNR over ASSR duration

In most applications, ASSRs are typically employed to determine the presence of a response, requiring that the response is above the significance threshold, with lesser emphasis on SNR differences between two ASSR responses. Therefore, we used a linear-mixed effect model to determine the duration required for each stimulus to reach the SNR threshold. We applied Hotelling’s *T*^2^ statistics to varying amounts of data for each modulation frequency, ASSR condition, and participant to achieve this goal. The data was sampled in one-minute intervals, ranging from one to eight minutes. We chose to fit a square root linear mixed-effects (LME) model to this data as a previous study has shown that the square root function best describes signal-to-noise ratio over increasing amounts of data (Elberling & Don, 1984). We initially fitted a linear mixed-effects (LME) model to our data, including all predictor variables and their interactions. However, as noted, like the other LME model, the interaction effects between the AM frequencies and conditions were insignificant. They were subsequently removed from the model, as shown in Equation 5. Like our previous model, the residuals of our second linear-mixed effect model do not follow a normal distribution, potentially resulting in reduced model power. Nevertheless, we again proceeded with the analysis as planned. In the subsequent analysis step, we employed the lower bound of the 95% prediction interval as an indicator to define the required duration for each stimulus to reach the SNR threshold. When the lower bound crosses the statistical threshold, it signifies the duration at which a new value will have a 95% probability of being statistically significant. Given the limited 8 minutes of data per stimulus, we utilized the LME model to make predictions across increasing amounts of data, using a step size of 6 seconds. This process continued until we identified the threshold where the lower bound crossed the significant threshold.

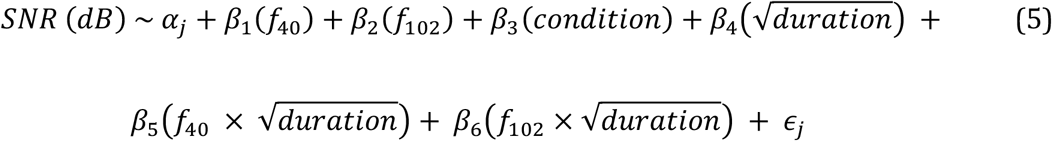

## Results

### Median SNR above significance threshold for each modulation frequency

The SNR of the evoked responses was calculated for each participant in the study using the eight-minute EEG recording. At eight minutes of data, the calculated F-statistic (F (2,479)) corresponding to an alpha of 0.05 is 3.01, which translates to 4.78 SNR dB (Dobie & Wilson, 1996). Across participants, the median SNR for single and multiplexed ASSR conditions was above 4.78 SNR dB, see Figure 4A. For the 40 Hz AM, all evoked responses are above the significance threshold. For the 3 Hz AM, 68% (15 out of 22) of the participants displayed significant responses to the multiplexed AM stimulus, while 81% (18 out of 22) displayed significant responses to the single AM stimulus. A similar pattern was observed for the 102 Hz AM, with 77% (17 out of 22) of participants displaying significant responses to the multiplexed AM stimulus and 91% (20 out of 22) displaying significant responses to the single AM stimulus. These results indicate that the single and multiplexed AM stimuli elicit significant synchronized activity in the EEG signal.

**Figure 4.**
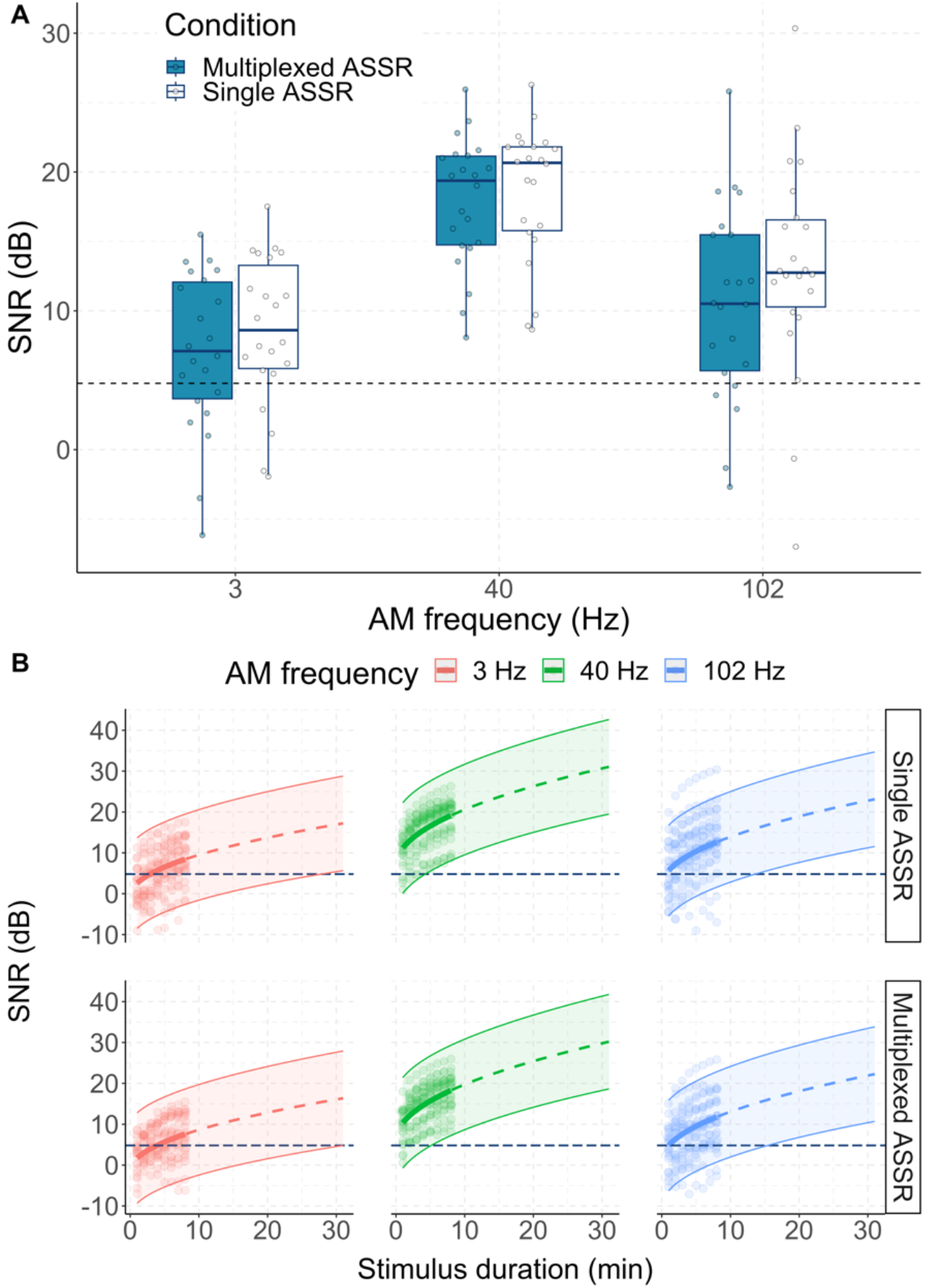
(A) The boxplots compare the amplitudes evoked by the single amplitude-modulation (AM) stimuli against those evoked by the multiplexed AM stimulus. A dot in each condition represents each subject. The solid line denotes the model fit based on the observed data, while the dashed line illustrates the model predictions for unseen data. (B) Compares the average signal-to-noise ratios (SNRs) evoked by the 3, 40, and 102 Hz AM frequencies for the single and multiplexed ASSR across stimuli duration. Each dot represents the computed SNR for each subject at each duration, while the bold lines show the average fit across subjects. The confidence interval of the model fit is depicted between the two outer lines.

Additional support for these findings comes from examining the frequency spectrum of the EEG. Across subjects, both stimuli exhibit distinct responses at each AM frequency for both single and multiplexed conditions, as depicted in Figure 5. Even at the individual subject level, the frequency spectra remain comparable between single and multiplexed AM stimulation. For instance, Subject S02 exhibits a pronounced evoked ASSR at the AM frequencies for multiplexed and single AM stimuli. In contrast, the responses of Subject S11 show lower amplitude, with only the peak at 40 Hz discernible.

**Figure 5.**
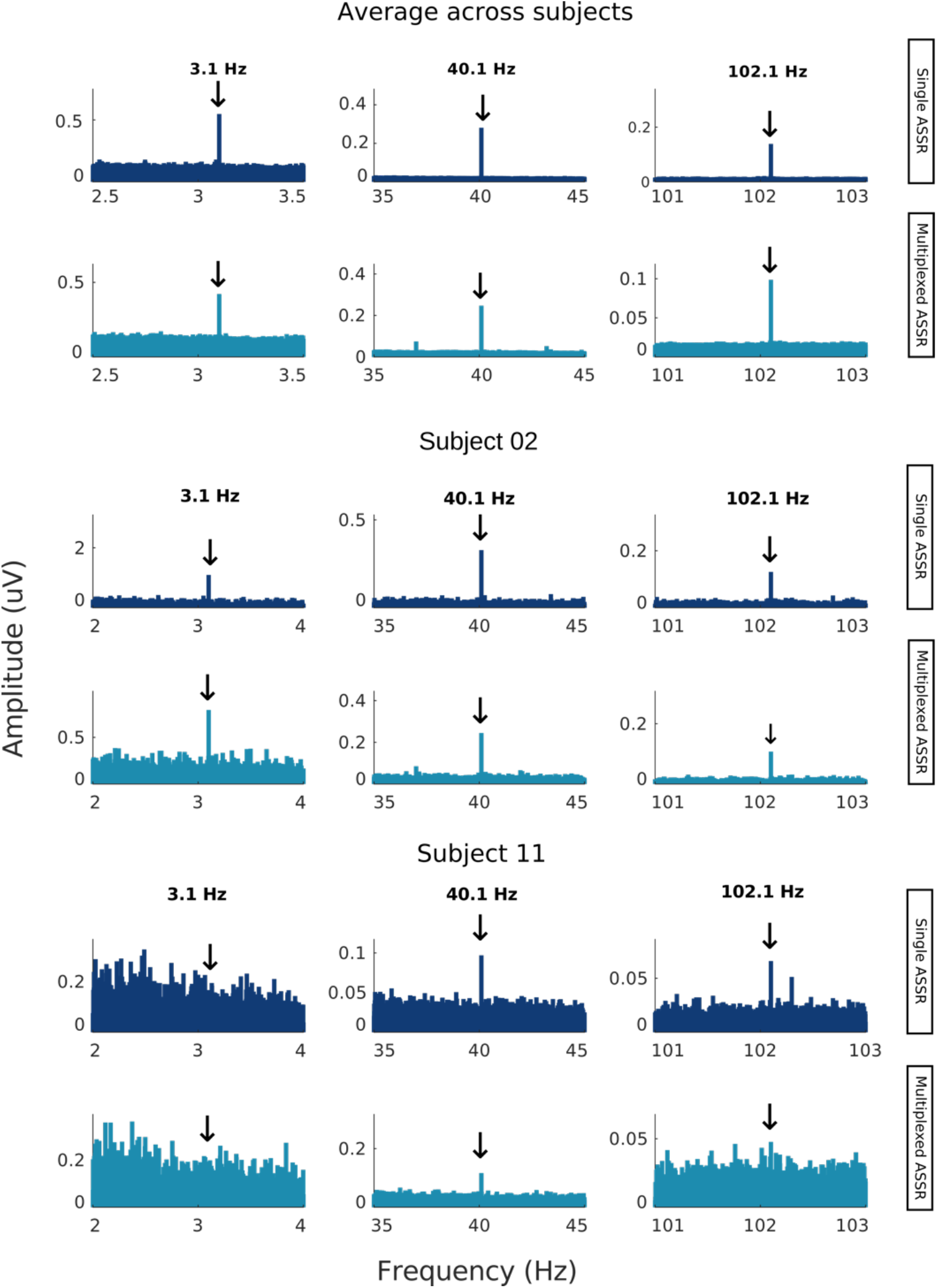
The figure shows the amplitude across frequencies of the ASSR response for the average across subjects and subjects S02 and S11, with arrows pointing to the amplitude-modulated frequencies of 3 Hz, 40 Hz, and 102 Hz.

### Significant difference in SNR for multiplexed vs. single AM stimuli

We investigated the potential difference in SNRs between the ASSRs evoked by a single AM stimulus and those evoked by a multiplexed AM stimulus.

Our LME model, yielding an *R*^2^ of 0.63, identified significant effects for the 40 Hz (*t* [110] = 10.91, *p* < .001) and 102 Hz (*t*[110] = 4.76, *p* < .001) categorical variables, as well as the intercept (*t*[49.86] = 7.65, *p* < .001). Additionally, the multiplexed stimulus significantly influenced the SNR value (*t*[110] = −2.08, *p* < .05). To ease the interpretations, we also conducted post hoc tests on the LME model; see Table 1. The pairwise comparison between the estimated marginal means of the single and multiplexed stimuli showed no significant difference in the SNR values (*difference* = 1.61 *SNR*(*dB*), *t*[113] = 2.05, *p* − *value* = 0.32), indicating that using a multiplexed AM stimulus does not significantly change the SNR values compared to using a single AM.

**Table 1:** displays the post hoc estimates for the Signal-to-Noise Ratio (SNR) and the corresponding 95% confidence interval (CI) obtained from the fitted LME model; see Equation 5. The estimates in the table represent the estimated marginal means while holding all other variables in the LME model constant.

| Multiplexed | AM frequency | Mean (SNR dB) | CI 2.5% | CI 97.5 % |
| --- | --- | --- | --- | --- |
| 0 | 3 | 8.64 | 6.33 | 10.95 |
| 1 | 3 | 7.03 | 4.72 | 9.34 |
| 0 | 40 | 19.02 | 16.70 | 21.33 |
| 1 | 40 | 17.40 | 15.09 | 19.71 |
| 0 | 102 | 12.61 | 10.30 | 14.92 |
| 1 | 102 | 11.00 | 8.69 | 13.31 |

### Similarity in sensor space patterns between multiplexed and single AM stimuli

Examining the sensor space across 64 channels, the amplitude patterns for each AM frequency exhibit remarkable similarity between the single and multiplexed stimuli, as illustrated in Figure 6A. Additionally, the similarity remains when we look at the SNR patterns within the sensor space, but it is slightly less pronounced at 102 Hz, shown in Figure 6B.

**Figure 6.**
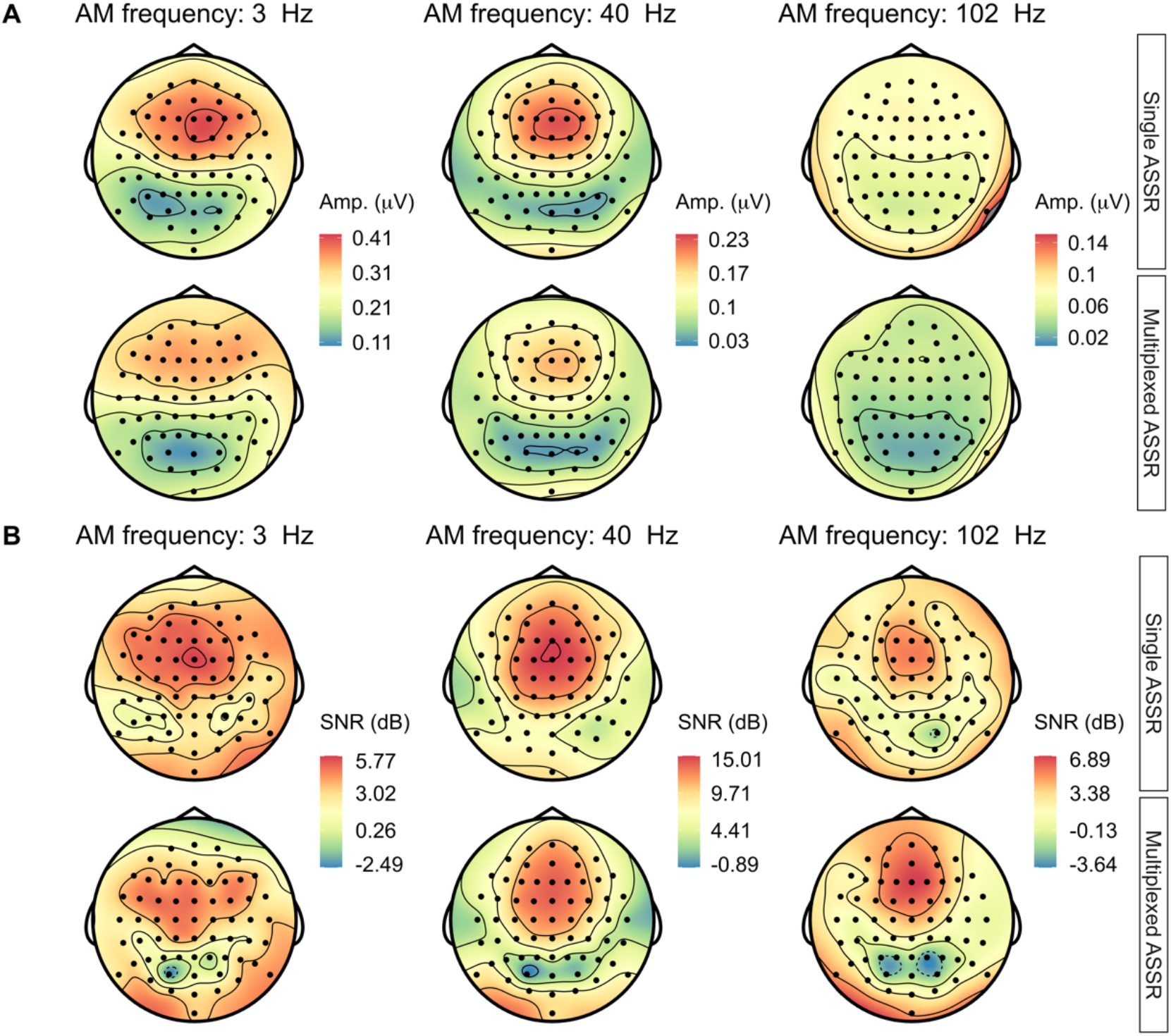
Topographic maps spanning all 64 channels illustrate the sensor space characteristics, showcasing (A) amplitude measures and (B) signal-to-noise ratio (SNR) (dB) across each amplitude-modulated (AM) frequency for both the single and multiplexed stimuli.

### Using a multiplexed AM stimulus reduces the recording time compared to using sequentially presented single AM stimuli

To complement our current findings, we sought to determine the minimum stimulus duration required to elicit a significant response for each type of auditory ASSR across all participants. Our fitted model, shown in Equation 5, had an *R*^2^ value of 0.71 and revealed the following fixed effects to be significant: ASSR condition (*t*[1034] = −3.59, *p* < .001), 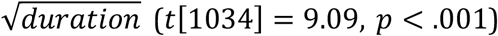, and 40 AM frequency (*t*[1034] = 7.07, *p* < .001), and 102 AM frequency (*t*[1034] = 2.33, *p* < .05). We also found that the interaction effect between the 40 Hz and 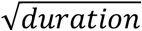 was significant (*t*[1034] = 2.28, *p* < .05). The post hoc test showed a 0.86 SNR dB (95% *CI* = [0. 39, 1.33]) difference between the multiplexed and single ASSR. Overall, both ASSR conditions showed a similar evolution of SNR across time, as depicted in Figure 4B. This evolution of SNR across time is seems to be driven by a reduction of noise and not by increasing amplitudes across time, see Appendix A.

We then investigated the minimum stimulus duration necessary for the lower bound of the 95% prediction interval associated with each AM frequency to cross the significance threshold. Our results revealed that, for the 3 Hz single AM frequency, 27 minutes and 48 seconds were necessary, 4 minutes and 18 seconds were required for the 40 Hz single AM frequency, and 13 minutes and 36 seconds were needed for the 102 Hz single AM frequency. In the case of the multiplexed AM stimulus, it took 31 minutes for the 3 Hz AM frequency, 5 minutes and 12 seconds for the 40 Hz AM frequency, and 15 minutes and 24 seconds for the 102 Hz AM frequency. The critical difference between the multiplexed and single stimuli was that the multiplexed stimulus allowed for the presentation of all AM frequencies simultaneously, resulting in a shorter overall duration of 31 minutes. In contrast, for the sequentially presented single stimuli, the stimulus duration for each AM frequency must be added together, resulting in a total duration of 45 minutes and 42 seconds. Our findings suggest that the multiplexed stimulus is more efficient and effective for eliciting significant ASSR responses across multiple AM frequencies.

## Discussion

This study compared the standard single AM stimulus, where each AM frequency is presented sequentially, to a novel approach called the multiplexed AM stimulus. The multiplexed AM stimulus is a single carrier wave modulated by multiple frequencies simultaneously, aiming to obtain information from various neural generators efficiently or simply measure the effect of modulation frequency. This approach contrasts with “multiple ASSRs,” where multiple carrier frequencies are used simultaneously. This study is the first to examine the ASSR to multiple simultaneously presented AM frequencies on a single carrier, specifically 3 Hz, 40 Hz, and 102 Hz.

We found a significant difference of 1.61 SNR dB between the single and multiplexed conditions on the SNR of the evoked response. Additionally, when we studied the effect of stimulus duration on the ASSR, where we tested durations ranging from one to eight minutes and calculated the SNR of the ASSR for each duration, we found that the single AM stimulus had a significantly 0.86 SNR dB higher than the multiplexed AM stimulus. However, it is worth noting that both the model and its residuals exhibit left skewness, increasing the likelihood of type I errors.

As a result, caution should be exercised in interpreting this significant difference as it may be a false positive result. If this finding is true, it would mean that considering a single frequency, single ASSRs are slightly more efficient. This result makes sense because the single sinusoidal modulators have been 100% amplitude modulated. When these modulators are combined into a multiplexed modulator, only 40.52% of its original amplitude is kept. These attenuations, although inconsistent across periods, remains consistent between AM frequencies in the frequency spectrum, as can be seen in Figure 1.

On average, the multiplexed AM stimulus took 31 minutes for the lower bound of the 95% prediction interval to cross the significance threshold across all three frequencies. In contrast, the single AM stimuli took 45 minutes and 42 seconds. Consequently, adopting the multiplexed AM stimulus is a valid alternative to using single AM stimuli, offering the advantage of reduced recording time. Over time, the amplitudes remain stable, illustrated in Appendix A, reinforcing the idea that the noise diminishes with prolonged stimulus duration, thus increasing the SNR.

Similar to previous studies, the SNRs of the ASSR exhibit an inverted U-shaped pattern across AM frequencies, with the highest SNR values appearing at around 40 Hz (Gransier et al., 2017, 2021). Previous studies have shown that the auditory system has the highest sensitivity to AM frequencies around 40 Hz and that this response originates from various neural generators (Farahani et al., 2017, 2020; Herdman et al., 2002; Johnson et al., 1988; Reyes et al., 2004; Weisz & Lithari, 2017). Furthermore, the percentage of significant ASSRs evoked by multiplexed and single AM stimuli are comparable to those reported by (Gransier et al., 2017, 2021) with all ASSRs evoked by the 40 Hz stimuli being significant. Notably, we found a higher number of significant responses for both the 3 Hz and 102 Hz AM frequencies, which can be attributed to the diotic stimulation used in our study, in contrast to the monaural stimulation employed by Gransier et al. (2017). Moreover, the stimuli in our study were presented at a sound intensity of 80 dB instead of 70 dB, which can also affect the amplitude of the ASSRs (Ménard et al., 2008).

The main result of our study shows that using a multiplexed AM stimulus can reduce recording time, which may be more practical for subjects prone to tiredness, loss of attention, and low arousal.

Another advantage of the multiplexed AM stimulus is the ability to simultaneously present a wide range of AM frequencies. This approach potentially enables the stimulus to target multiple neural generators in the auditory pathway. Higher AM frequencies above 80 Hz tend to elicit more subcortical activity, primarily from the brainstem (Bidelman, 2015; Farahani et al., 2020; Herdman et al., 2002; Luke et al., 2017). In contrast, lower AM frequencies below 20 Hz elicit more cortical activity, primarily from the auditory cortex 5/21/24 11:52:00 AM. However, a challenge inherent in all ASSR stimuli is associating modulation frequency ranges with *precise* neural generators, particularly in pinpointing the exact contribution of each neural generator to the ASSR response. In addition, the multiplexed stimulus yields a more complex modulation spectrum. It is highly non-linear, thus making it possible that there are unexpected interactions between the ASSR responses. Despite these challenges, our findings indicate no noticeable differences in SNRs and amplitudes between the multiplexed ASSR and the single ASSR. The topographic maps also show an absence of distinct differences between both ASSRs, which at least does challenge the notion that either brain or cortical generators predominantly drive the multiplexed ASSR. Furthermore, the 40 Hz AM frequency amplitude topographies closely resemble those reported in other studies (Johnson et al., 1988; Neklyudova et al., 2021; Saupe et al., 2009).

Based on the study results, a potential avenue for future research would be to improve the multiplexed AM stimulus by increasing the number of AM frequencies included. This approach would allow for an analysis of the apparent latency (Picton et al., 2003), which can provide more information about the neural generators involved in the multiplexed ASSR and therefore provide greater insights into the integrity of the auditory pathway (L. Wang et al., 2021). In this study, we used multiplication to create the multiplexed stimulus, yielding a more complex modulation spectrum. However, for future research, choosing addition over multiplication will facilitate the development of a simple spectrum, thereby reducing potential interacting and confounding factors.

Another avenue might investigate multiplexed sweeps, where sweeps on lower frequency ranges are simultaneously combined with sweeps on higher frequency ranges; this might help to better capture ASSRs from each neural generator, as the specific AM frequency that neural generators respond best to can vary strongly across individuals (Gransier et al., 2021).

## Conclusion

In conclusion, our study revealed that employing a multiplexed ASSR is more time-efficient than using a sequentially presented single ASSR.

## Appendix

**Appendix A:**
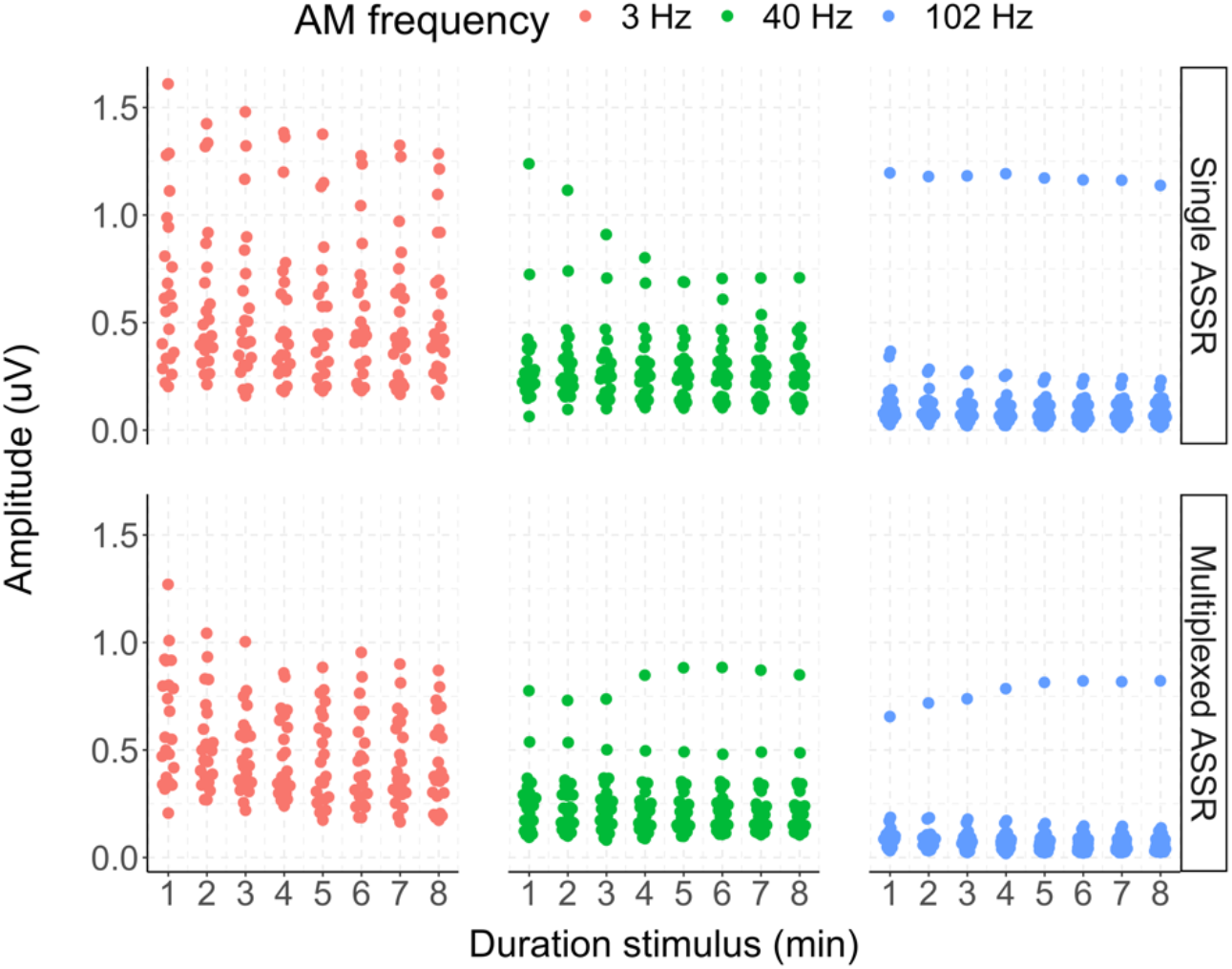
Compares the amplitudes evoked by the 3, 40, and 102 Hz amplitude-modulated (AM) frequencies for the single and multiplexed auditory steady-state response (ASSR) across stimuli duration. Each dot point represents the amplitude for an individual subject at each specified duration

